# Amplitude and timescale of metacommunity trait-lag response to climate change

**DOI:** 10.1101/850289

**Authors:** Jon Norberg, Helen Moor

## Abstract

Climate change is altering the structure and functioning of communities ^1^. Trait-based approaches are powerful predictive tools that allow consideration of changes in structure and functioning simultaneously ^2, 3^. The realised biomass-weighted trait distribution of a community rests on the ecophysiology of individuals, but integrates local species interactions and spatial dynamics that feed back to ecosystem functioning. Consider a response trait that determines species performance (e.g. growth rate) as a function of an environmental variable (e.g. temperature). The change in this response trait’s distribution following directional environmental change integrates all factors contributing to the community’s response and directly reflects the community’s response capacity ^3^.

Here we introduce the average regional community trait-lag (TL_MC_) as a novel measure of whole-metacommunity response to warming. We show that functional compensation (shifts in resident species relative abundances) confers initial response capacity to communities by reducing and delaying the initial development of a trait-lag. Metacommunity adaptive capacity in the long-term, however, was dependent on dispersal and species tracking of their climate niche by incremental traversal of the landscape. With increasing inter-patch distances, network properties of the functional connectivity network became increasingly more important, and may guide prioritisation of habitat for conservation.

---

Anticipating climate change effects on the biosphere requires a process-based/mechanistic understanding of community reorganisation in response to changing conditions. Commonly used approaches based on species distribution modelling often assume universal dispersal ^4^, thereby ignoring the essential patchiness of most habitats as well as habitat fragmentation caused by human land use ^5, 6^. Niche/Distribution modelling further tends to assume that species move in isolation, when in reality the landscape is filled with potential competitors that can resist colonisation or drive vulnerable species to extinction ^4^. Lastly, studies of ecological responses to climate change tend to focus on individual organisms or species, and rarely assess whole community and ecosystem responses ^1^.

Trait-based approaches facilitate the integration of responses of individual organisms with species interactions as well as the scaling to community and ecosystem level changes. Describing communities in terms of the abundance- or biomass-weighted distribution of relevant traits can simultaneously integrate the factors affecting community structure and assess potential effects on ecosystem functioning. Community trait distributions characterised by the four central moments (mean, variance, skewness, and kurtosis) hold more information than species richness and evenness alone ^3^, and integrate functional diversity components ^7^.

The variance of the community distribution of a response trait that affects performance (e.g. growth) represents the response diversity ^8, 9^ that directly determines the capacity of the community to adapt to directional environmental change ^10^. In fact, given the general growth function in relation to the driver and the biomass weighted distribution of the response trait(s) in the community, it should be possible to predict the response capacity of the whole community ^11^. Specifically, a change in environmental conditions will cause the mean of the trait distribution to shift as species better adapted to new conditions increase in relative abundance (Fig 1), thus maintaining community productivity through functional compensation ^12^. Recent theory ^3, 10, 13^ predicts the mean to lag behind the novel optimum conditions at a magnitude that depends on the rate of environmental change and the variance of the trait distribution ^3^. The lag of the trait distribution mean behind current conditions (hereafter trait-lag, TL) represents the realised momentary response of the community. Importantly, a greater TL coincides with relatively lower community net productivity, as the average of species growth rates remains below optimal. This holds for any response trait affecting growth. For performance traits affecting other fitness components ^14^, it would imply lower survival rates or lower reproductive output.

**Figure 1:**
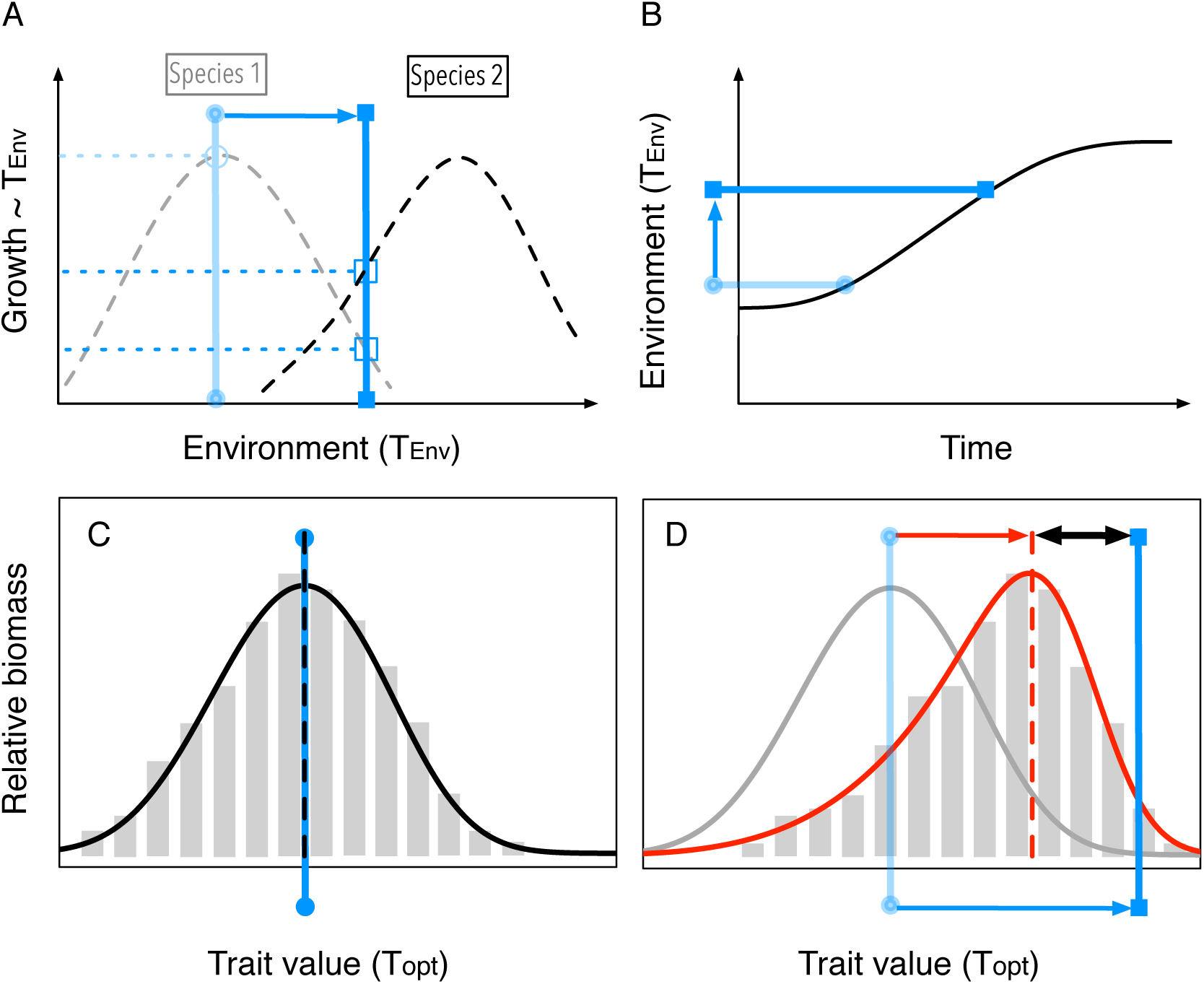
Illustration of the trait-lag (TL) of a biomass-weighted trait distribution of a community. **A:** Performance/growth curves in response to temperature of two species with different temperature optima. Blue bars and lines indicate their respective growth rates in two environments differing in temperature. **B:** Temperature increase due to climate change. **C:** During stable conditions, the community trait distribution converges around the current environmental optimum, indicated by the blue handle with round ends. The community weighted mean trait (CWMT) coincides with the current optimum trait value. Grey bars indicate relative biomasses of species with particular trait values of the response trait in question, here the temperature optimum T_opt_. **D:** Consider a change in environmental conditions as in panel B, indicated by the blue arrow, that shifts the current optimum to higher values of T_opt_. The new optimum is indicated by the blue handle with square ends. Due to the increase in abundance of species with trait values that match the new environment more closely, the mean of the trait distribution has shifted as well, tracking current conditions (indicated by the red arrow). Tracking has not been complete, however, and the CWMT has developed a lag, the trait-lag TL, behind the new optimum (indicated by the black two-headed arrow).

In a metacommunity situation, the dependence of TL on trait variance is complicated by the interplay of local and regional dynamics. The local variance here is affected by both local species interactions and short term compensatory dynamics in response to environmental fluctuations (e.g. stronger competition decreases the number of coexisting species and hence the variance, while environmental noise maintains a greater variance) as well as regional processes (immigration of species with different trait values can maintain or increase the trait variance).

With a focus on regional aggregate properties, the average TL across all communities in the region can be used as a quantitative measure of the momentary whole-metacommunity response to changing conditions. The temporal dynamics of this regional trait-lag TL_MC_ yield a direct measure of metacommunity response capacity.

Here we used a spatially explicit metacommunity model to explore changes in community distributions of a temperature optimum trait in response to climate-change driven warming. The temperature optimum trait determines species fitness (growth rate) as a function of current environmental conditions (temperature as ‘performance filter’ ^11^). Local coexistence is maintained by both weaker interspecific compared to intraspecific competition ^15^ and (weakly autocorrelated) environmental noise that causes local compensatory dynamics between species ^12^.

The model tracks the lag of the community weighted mean trait (CWMT) behind shifting optimal conditions averaged across all patches in the metacommunity (TL_MC_). This time dynamic of the regional lag TL_MC_(t) was characterised with regard to timescales of development (t_Dev_) and recovery (t_Rec_), as well as the amplitude TL_Max_ of the lag. The analysis focuses on a novel measure to quantify the response capacity of the whole region: The integral over time of the regional average lag ∫TL_MC_ is a single value metric that captures whole-metacommunity response to warming; we propose the use of its inverse 1/ ∫TL_MC_ as a measure of metacommunity response capacity. This response capacity was compared for metacommunities with different species/functional group characteristics (strength of interspecific competition and population growth rates) and landscape characteristics (inter-patch distance and patch arrangement, quantified as topology metrics of the connectivity network). Additionally, TL_MC_(t) could be fully determined by decomposition into contributions from local dynamics (shifts in relative abundance of competing resident species in the absence of dispersal, or local species sorting TL_SS_) and contributions from regional dynamics (immigration from dispersal, TL_Disp_, and, as part of dispersal, the incremental traversal of the landscape by establishment in stepping stone patches, TL_Trav_).

## Results

### TL_MC_ over Time

All metacommunities developed a trait-lag TL_MC_ in the wake of climate change. TL_MC_ always reached a maximum **TL**_**Max**_ and subsequently decreased again as the metacommunity started to recover and the developed trait-lag diminished (Fig 2 A). TL_Max_ was negatively correlated with response capacity (1/ ∫TL_MC_), and linearly increased with inter-patch distance (hereafter distance). Corresponding to higher TL_Max_ at greater distance, the **initial development of the lag** continued longer at greater distances, i.e. **t**_**Dev**_ increased with distance and TL_Max_ was reached later. **Time to recovery** back to half of TL_Max_ increased with distance up to the greatest distance (16 km), where **t**_**Rec**_ was not reached and a large trait-lag persisted throughout the entire simulation (runtime 3000 yrs), long after warming had ceased and environmental conditions had stabilised.All three aspects of the temporal dynamic of TL_MC_ (TL_Max_, t_Dev_, and t_Rec_) were positively correlated and responded in the same way as response capacity to levels of competition and growth (Fig 2), with the exception of the largest landscape with greatest inter-patch distances (Suppl Fig S1). That is to say, metacommunities with the highest response capacity had the lowest lag amplitude TL_Max_, developed a lag faster and recovered earlier.

**Figure 2:**
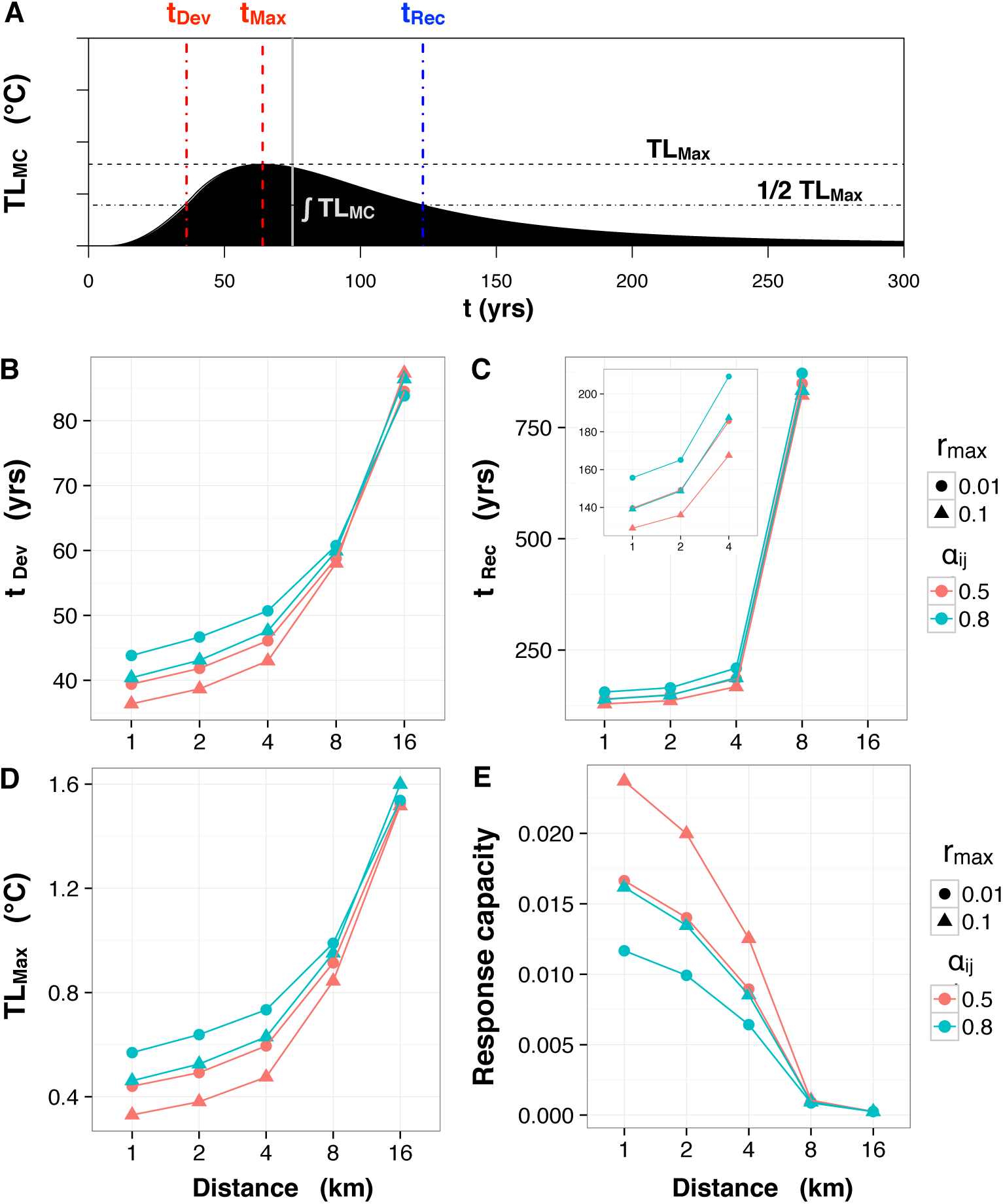
Example of a time dynamic of the metacommunity trait-lag TL_MC_ and mean responses of its aspects to factor levels. **A:** The time dynamic of TL_MC_ was characterised by its amplitude TL_Max_ and timescales of development (t_Dev_, dot-dashed red line) and recovery (t_Rec_, dot-dashed blue line). The full grey line indicates the half-time of the climate change warming curve. TL_MC_ was summarised by its integral over time ∫TL_MC_ (black area). The inverse of the integral 1/ ∫TL_MC_ was used a a measure of metacommunity response capacity. **B-E:** Mean responses of aspects of the TL_MC_) time dynamic and response capacity to factors inter-patch distance (x-axis), maximum growth rate r_max_ (circle= 0.01; triangle= 0.1), and strength of interspecific competition *α*_*ij*_ (red= 0.5; blue = 0.8). **B:** Half-time of lag development t_Dev_. **C:** Time of recovery to half of TL_Max_, t_Rec_ (not reached at largest extent; the insets shows a close-up for the three shortest distances.). **D:** Maximum trait-lag TL_Max_. **E:** Response capacity 1/ ∫TL_MC_. All aspects of TL_MC_ (t_Dev_, t_Max_ (not shown), t_Rec_, and TL_Max_) were positively correlated.

### ∫TL_MC_ and Response Capacity The integrated regional lag ∫TL_MC_

(corresponding to the black area in Fig 2) monotonically increased with inter-patch distance, highlighting the importance of dispersal in reducing TL_MC_. At the two greatest distances, 8 and 16 km, recovery of TL_MC_ was slow, and ∫TL_MC_ increased exponentially.

The inverse of the integrated regional response (1/ ∫TL_MC_) was used as a single-value metric of whole-metacommunity **response capacity** and analysed with regard to factor manipulations (Fig 2 E). Inter-patch distance had the largest effect on response capacity (Table 1). Response capacity steadily decreased with increasing distances, as dispersal became increasingly less efficient and rescue and re-organisation of local communities ultimately failed (much reduced recovery of the regional trait-lag in the largest landscape).

**Table 1:** Anova results of main effects and interactions of factors distance, competition (*α*_*ij*_), maximum growth rate (*r*_*max*_) and network combination (Network) on metacommunity response capacity (= 1/ ∫ *TL*_*MC*_). The intention is not to examine statistical significance but to indicate the relative effect sizes of factors, calculated as *η*^2^. Commonly used benchmarks for small, medium and large effect sizes are 0.02, 0.13, 0.26 respectively (Cohen 1988, Miles and Shevlin 2001).

| | Sum Sq | Df | F value | $\eta^2$ |
| --- | --- | --- | --- | --- |
| Distance | 0.00 | 4 | 3199.2 | 0.995 |
| $\alpha_{ij}$ | 0.00 | 1 | 708.7 | 0.905 |
| $r_{max}$ | 0.00 | 1 | 1592.5 | 0.886 |
| Network Combination | 0.00 | 7 | 2.7 | 0.034 |
| Distance : $\alpha_{ij}$ | 0.00 | 4 | 148.3 | 0.877 |
| Distance : $r_{max}$ | 0.00 | 4 | 334.4 | 0.856 |
| $\alpha_{ij}$ : $r_{max}$ | 0.00 | 1 | 106.2 | 0.312 |
| Distance : Network | 0.00 | 28 | 1.2 | 0.064 |
| $\alpha_{ij}$ : Network | 0.00 | 7 | 0.1 | 0.001 |
| $r_{max}$ : Network | 0.00 | 7 | 7.4 | 0.007 |
| Distance : $\alpha_{ij}$ : $r_{max}$ | 0.00 | 4 | 24.3 | 0.246 |
| Distance : $\alpha_{ij}$ : Network | 0.00 | 28 | 0.1 | 0.005 |
| Distance : $r_{max}$ : Network | 0.00 | 28 | 1.8 | 0.027 |
| $\alpha_{ij}$ : $r_{max}$ : Network | 0.00 | 7 | 0.7 | 0.001 |
| Distance : $\alpha_{ij}$ : $r_{max}$ : Network | 0.00 | 28 | 0.2 | 0.002 |
| Residuals | 0.00 | 2239 |  |  |

Both the strength of interspecific competition *α*_*ij*_ and maximum growth rate *r*_*max*_ affected response capacity, but relevant mechanisms differed between patch distance levels (interaction terms, Table 1). Both the effect of competition and of *r*_*max*_ decreased with increasing distances. Overall, communities (or functional groups) characterised by weaker interspecific competition and faster growth rates had the highest response capacity, while communities with strong competition and slow growth rates had the lowest (Fig 2 E). In terms of life history theory or the CSR strategies of Grime ^16^, ruderals or opportunistic species would have greater response capacity, while competitive but slow growing species would exhibit lower response capacity. Note however that neither fecundity nor mortality or lifespan were different between these groups here, such that differences in responses were only due to growth and competitive interactions. Combinations of strong competition with fast growth, and weak competition with slow growth showed intermediate response capacity and behaved similarly, suggesting that growth and competition had opposite and countervailing effects. *r*_*max*_ had a slightly larger effect at weaker competition levels. The 3way interaction between distance, *α*_*ij*_ and *r*_*max*_ was due to a shift in the interaction at the greatest distance (16 km) (Suppl Fig S1).

Network properties that characterised the functional connectivity of the landscape had overall comparatively small effects on response capacity, but their effect became noticeable at the two greatest distance levels (8 and 16 km) (interaction term, Table 1). The best combination differed between distance levels (Suppl Fig S2). At both distance levels (8 and 16 km), combinations with a higher degree of clustering and a shorter characteristic path length corresponded to higher response capacity. Interestingly, at the greatest inter-patch distances (16 km), a larger leading eigenvalue of the connectivity matrix appeared to confer higher response capacity (Fig 3).

**Figure 3:**
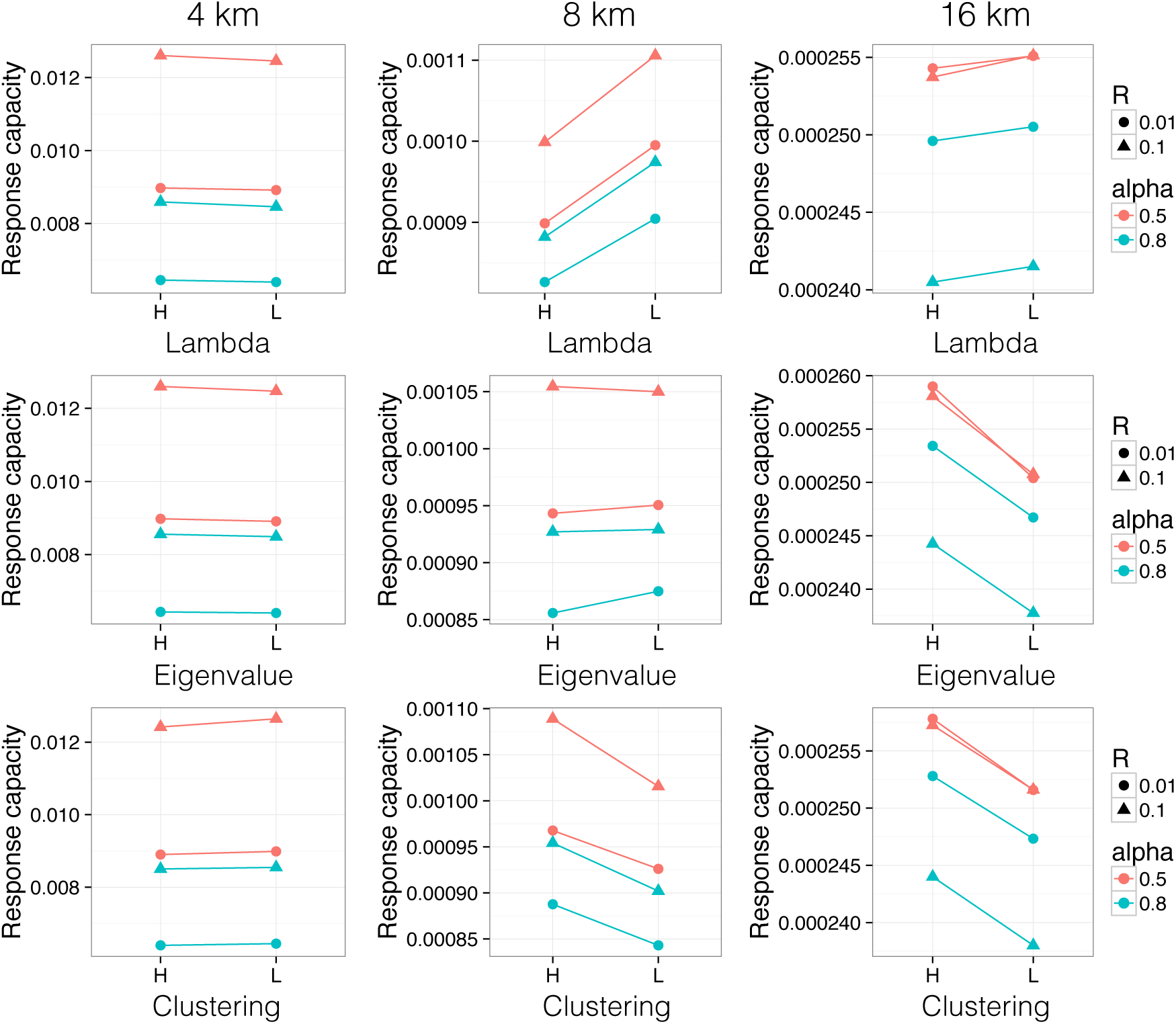
Comparison of mean response capacity between high (H) and low (L) values of network properties characteristic path length (Lambda), leading eigenvalue (Eigenvalue) and clustering (Clustering) at the three largest inter-patch distance levels. In landscapes of distances up to and including 4 km, network properties showed no correlation with response capacity. At the two largest patch distances, high clustering corresponded to higher response capacity, while lamdba and eigenvalue affected response capacity differentially at distance 8 km compared to 16 km.

Overall, at inter-patch distances up to 4 km, dispersal appeared to be efficient and differences in response capacity were influenced more by local dynamics and effects of growth and competition. At distance level 8 km, dispersal became limiting, and the effect of local community dynamics less important. The largest landscapes with the greatest inter-patch distance (16 km) behaved qualitatively differently.

### Local and Regional Components

The realised regional response that mediated TL_MC_ could be decomposed into local and regional response mechanisms or components, the relative importance and timescale of which depended on manipulated factors (above all inter-patch distance, Fig 4). After the onset of climate change, shifts in the relative abundances of local resident species present, i.e. local species sorting **TL**_**SS**_ (corresponding to the orange area in Fig 4), consistently and strongly contributed to buffering the initial development of TL_MC_ in response to warming. Note that TL_Max_ in all cases remained substantially lower than the maximum possible lag: with no change in species abundances and trait distributions, TL_Max_ would ultimately have corresponded to the maximum of environmental temperature change (5 °C). Local response diversity thus was high enough to delay the development of a local community trait-lag, and to substantially reduce TL_MC_ relative to he maximum possible lag given by the shift in the environmental optimum (the warming curve; top black line in Fig 4). At all extents, TL_SS_ accounted for >95% of lag reduction at t_Dev_ (Suppl Fig S3). Local shifts in resident species abundances determined initial response capacity.

**Figure 4:**
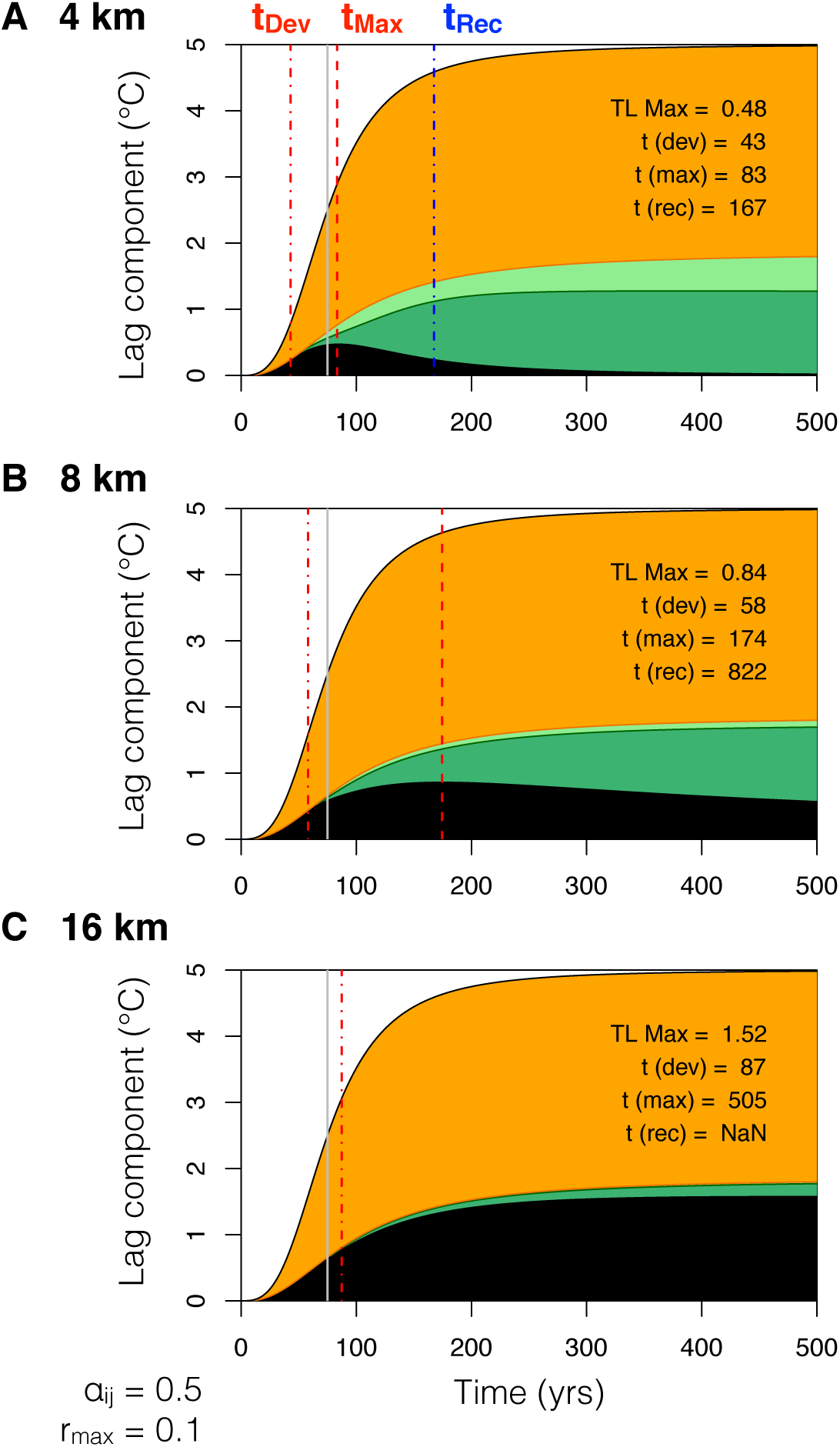
Time dynamic of the regional trait-lag TL_MC_ (black area) for 500 yrs after the onset of climate change (CC) at time = 0 (total runtime 3000 yrs) for the three largest inter-patch distances (A: 4, B: 8, C: 16 km). Coloured areas show the contributions of components to reducing the lag relative to climate change induced warming (i.e. the maximum possible lag, top black curve): local shifts in abundances of resident species (TL_SS_, orange area), dispersal (TL_Disp_, light and dark green areas), and partial contribution to dispersal of incremental traversal of the landscape (TL_Trav_, dark green area). Vertical lines and text indicate time points of i) dot-dashed red: t_Dev_, ii) full grey: halftime of CC, iii) dashed red: t_Max_, and iv) dot-dashed blue: lag recovery halftime t_Rec_.(Shown are means across network combinations for *α*_*ij*_ = 0.5, *r*_*max*_ = 0.1.)

Already during climate change dispersal **TL**_**Disp**_ began to contribute to the attenuation of the lag via the immigration of species better adapted to the newly changed local conditions (light and dark green areas in Fig 4). By t_Max_, TL_Disp_ contributed with 8-16% to lag reduction (Suppl Fig S3). Lag development thus was slowed down and restrained by dispersal. The relative importance of dispersal increased from inter-patch distances 2 to 8 km, and dropped again at the largest distance level (dispersal limitation) (Suppl Fig S3 B). By t_Rec_, dispersal had increased its contribution to lag reduction to 25-32% (Suppl Fig S3 C). The relative contribution of dispersal now increased monotonically with distance. Dispersal thus fuelled regional re-organisation and trait-lag recovery.

With increasing inter-patch distances, an increasingly greater part of dispersal could be attributed to the incremental traversal of the landscape (**TL**_**Trav**_, dark green area in Fig 4) by species that could utilise intermediate patches as stepping stones, i.e. immigrate, establish, and disperse further across the landscape. The contribution of TL_Trav_ to lag reduction increased up to the greatest distance, where traversal became less efficient (Fig 4 C; Suppl Fig S3 E). TL_Trav_ became most important during recovery of TL_MC_ in later phases of climate change and after temperature change saturated. Species range shifts across the landscape thus temporally lagged behind environmental change and continued after climate change induced warming had come to a halt.

The timescales and amplitudes of the component functions shed light on the interacting mechanisms involved in the overall response (Fig 5). The lag developed in the absence of dispersal (TL_SS_) was independent of patch distance and landscape extent, but was affected by growth and competition levels (Fig 5 A). Dispersal dynamics on the other hand began to counter lag development earlier at smaller distances and the magnitude of lag reduction due to dispersal was much reduced at the largest distance level (Fig 5 B). The contribution of TL_Trav_ to lag reduction steadily increased with distance, except for the largest distance level, where range shifts across the landscape were rare (Fig 5 C).

**Figure 5:**
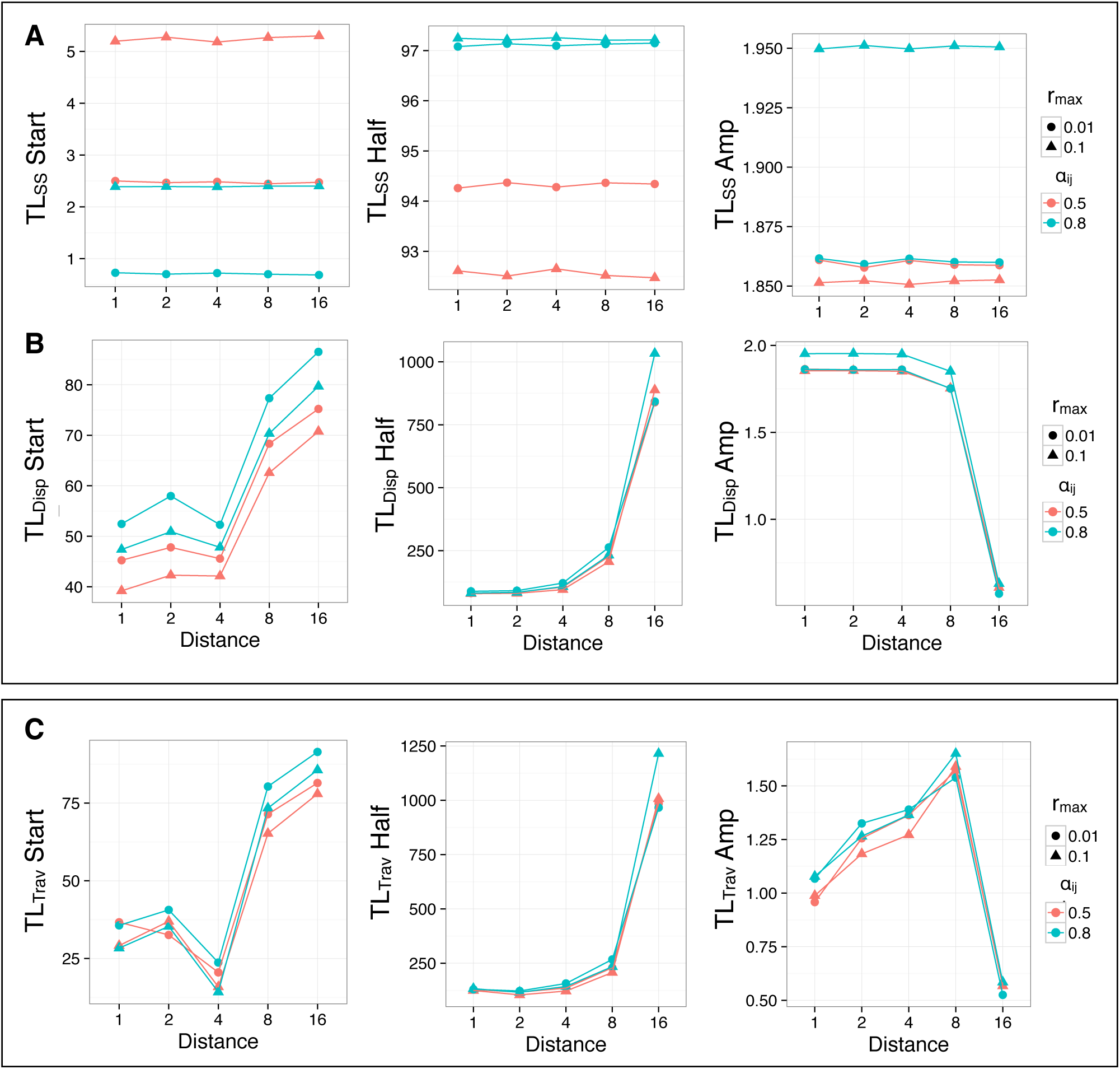
Fit estimates of three parameters (Start, Amp, Half) of the sigmoid functions fitted to components of the lag (cf. Methods, Eq 4). Shown are means for factor levels of patch distance (x-axis), maximum growth rate (symbol), and competition *α*_*ij*_ (colour). **A:** TL_SS_, and **B:** TL_Disp_, which together fully determine TL_MC_. **C:** TL_Trav_, the part of TL_Disp_ attributed to incremental traversal of the landscape. Note that TL_SS_ corresponds to the lag developed in the absence of dispersal, while TL_Disp_ corresponds to the *reduction* of this lag by dispersal. *TL*_*MC*_(*t*) = *TL*_*SS*_(*t*) − *TL*_*Disp*_(*t*).

Metacommunities with the highest response capacity (weak competition and fast growth, “fast ruderals”) were characterised by the latest initial development of a trait-lag TL_MC_, the lowest absolute trait-lag in the absence of dispersal, and the earliest onset of dispersal and traversal dynamics. Metacommunities with the lowest response capacity (strong competition and slow growth, “slow competitives”) showed the earliest initial development of TL_MC_ and dispersal started to contribute the latest. The timescales of component contributions thus were most important in determining overall response capacity. Weak competition and faster growth rates allowed the combination of high initial response capacity, i.e. a late and weak initial lag development, and the early onset of dispersal dynamics, which together conferred the highest response capacity. Strong competition and slow growth on the other hand lead to fast development of TL_MC_ in conjunction with a late onset of dispersal dynamics, which depressed overall response capacity.

Metacommunities with both strong competition and fast growth rates showed an interesting pattern that resulted in an intermediate response capacity: while the lag developed in the absence of dispersal was greatest, dispersal also contributed most to recovery of that lag (Suppl Fig S3). Due to competitive exclusion during stable conditions, these communities had the lowest initial trait distribution variances (Suppl Fig S4), which hampered the initial response capacity of local communities and depressed overall response capacity by causing a large initial lag. Eventhough dispersal and traversal of the landscape here contributed most to the recovery of the trait-lag during later phases of the response (Suppl Fig S3), this was insufficient to completely overcome the initially developed lag.

## Discussion

We have proposed to follow and examine the development and recovery of the average regional community trait-lag in response to climate change, and to use its time integral to derive a measure of whole-metacommunity response capacity. A focus on trait distributions and the trait-lag time dynamic has several advantages: i) the trait-lag can be conveniently summarised into a single measure of regional response capacity, ii) it allows to disentangle the mechanisms contributing to community re-organisation, iii) it yields information on the timescale of the dynamical response, and iv) it allows for predictions regarding concurrent changes in ecosystem functioning (i.e. rates of primary productivity: the smaller the trait-lag, the higher the maintenance of productivity rates).

### Response capacity depends on the interplay of local and regional dynamics

The dynamical response of the metacommunity to changing environmental conditions involved two phases that were characterised by different mechanisms. First, during the initial response phase, local shifts in relative abundances of resident species (functional compensation ^12^) conferred initial response capacity to communities by reducing and delaying the development of a trait-lag. Second, in a regional response and re-organisation phase the trait-lag development was slowed down and then reversed, and community functioning recovered. This buffering and recovery of the trait-lag depended on regional metacommunity re-organisation via species migration. Long-term adaptive capacity thus was only possible through dispersal and species tracking of their climate niche by incremental traversal of the landscape. Dispersal maintained local compensatory dynamics via a spatial insurance effect ^17^, where species adapted to warmer conditions could shift their ranges and compensate for the loss of species less well adapted to newly changed conditions. It was the combination of high initial response capacity with efficient dispersal that resulted in the highest overall response capacity.

### Determinants of initial response capacity

Local response capacity has previously been shown to be directly proportional to the variance of community response trait distributions ^10, 18^. This conjecture is the trait-based equivalent of the insurance hypothesis ^19^ or an analogue of the notion of response diversity ^9^, where a high variance implies higher levels of local coexistence of species with different environmental optima such that one species may compensate for the loss of another to maintain levels of ecosystem functioning under changing conditions (functional compensation ^12^).

In our model, trait variances were generally high. This resulted from both weaker inter-*versus* intra-specific competition as well as from the inter-annual variability in experienced temperatures that generated temporal fluctuations in conditions, both promoting local species coexistence ^15^. Variance then played a limited role and was only weakly related to initial response capacity. Stronger competition depressed initial community trait variance by competitive exclusion during stable climates, resulting in comparatively lower response diversity, an earlier lag development and a greater trait-lag in the absence of dispersal.

Faster growth rates on the other hand were overall beneficial for initial response capacity, in spite of associated lower community trait variances. Because local response diversity was sufficiently high in all cases, the speed of the response was the decisive determinant. Faster growth rates accelerated competitive displacement and species turnover rates, which allowed communities to track changing conditions faster and delayed the development of a trait-lag.

### Determinants of long-term adaptive capacity

Although local response capacity contributed substantially to absolute lag reduction (orange area in Fig 3), it was limited to an initial shift and a temporary tracking of temperature change by community trait distributions only. Regional dispersal dynamics were instrumental for the attenuation of lag development and community re-organisation that would reduce and recover the trait-lag. Directional environmental change such as regional warming requires dispersal-mediated range shifts of species to maintain community functioning.

Again, the speed of the response had the greatest effect on regional response capacity. The timing of the onset of dispersal dynamics (TL_Disp_ Start) determined how large the lag would get, and was the best indicator for lag recovery rates and thus overall response capacity. Earlier and faster dispersal dynamics curbed lag development, and corresponded to a greater absolute contribution of dispersal to lag reduction.

#### Effects of functional group characteristics

The earliest onset and the strongest contribution of dispersal dynamics occurred under weak competition levels combined with fast growth rates (“fast ruderals”). These conditions allowed immigrants with more optimal trait values to quickly establish and increase in biomass, facilitating range shifts via traversal of the landscape along the network of patches and resulting in the highest regional response capacity.

At the other extreme, metacommunities of species characterised by strong competitive interactions and slow growth (“slow competitives”) showed the weakest response capacity. Lag development took longest in these metacommunities, but they also developed the largest lag and recovered more slowly. This community inertia resulted from persistence of species with suboptimal trait values through preemptive dominance and from slow growth of colonisers, hampering competitive displacement. Dispersal contributed the least in these metacommunities, potentially due to competitive resistance from resident species, that, in conjunction with slow growth rates, impeded colonisation and range shifts by traversal. Slow changes in plant communities in response to changing conditions, along with the maintenance of productivity levels during the first phase of climate change, thus not necessarily imply high long-term response capacity. Our results suggest that such resistant but slow-turnover communities may develop the greatest long-term lag, with long-lasting effects on productivity levels. Persistence of ultimately maladapted species essentially creates a functioning debt ^20^ that continues long after climate change has slowed down. Because of their slow turnover rates, rescue from plastic differentiation into ecotypes or evolutionary dynamics ^21^ is unlikely to play a role. In spite of initial resistance, the functioning of such communities is thus most vulnerable in the long run.

#### Effects of landscape characteristics

The size of the landscape and resulting distances between patches were the strongest mediators of long-term response capacity. In metacommunities spread over larger landscapes lag development continued for longer, larger lag amplitudes were reached, and communities recovered more slowly. Landscape design aimed at realism but the number of patches was constrained by computation time. The solution was to keep the number of patches constant, while scaling the size of the landscape (and hence inter-patch distances) along with the temperature gradient spanned by the landscape (constant 1°C/100 km). Due to this model setup, the effect of distance is the result of three interacting effects.

First, increasing inter-patch distances decreased the likelihood and efficiency of dispersal, affecting both migration within the landscape as well as immigration from the wider region. Second, because the temperature gradient within the landscape was smaller at smaller distances, patches within the landscape were more similar in their experienced temperatures and the regional set of initially present species smaller. In small landscapes, adaptation to warming was thus dependent on immigration from the regional species pool, while in large landscapes, range shifts of species present in the landscape played a larger role. Third, temporal environmental variability was modelled as stochastic and weakly temporally autocorrelated noise in experienced temperatures. These fluctuations were the same throughout the landscape, i.e. patches fluctuated in synchrony. In larger landscapes, where patches differed more in baseline temperature, this increased niche diversity in both space and time. In smaller landscapes, however, fluctuations were large relative to baseline differences between patches, and thus contributed to overall synchronisation of the whole metacommunity, decreasing niche diversity and spatial insurance effects ^17^. This entanglement complicates the interpretation of observed patterns and explains why we do not observe a uni-modal relationship of response capacity with inter-patch distance, as would be expected for the relationship between dispersal rate and community diversity and stability ^17, 22^. The model would benefit from a setup that separates these effects (see Limitations).

### Connectivity network properties matter when habitat is scarce

In larger landscapes the potential of dispersal to reduce the trait-lag and to aid in the regional re-organisation and adaptation to new conditions was limited. Increasing inter-patch distances reduced dispersal efficiency and impeded recovery of the trait-lag, leading to persistent high lags in landscapes where habitat and stepping stones were scarce.

With increasing inter-patch distances and a wider temperature gradient in the landscape, the incremental traversal of the landscape via stepping stone patches became ever more important to metacommunity adaptive capacity. True range shifts were required to maintain whole-metacommunity functioning. In such landscapes, network properties of the functional connectivity network began to matter. Their effect was comparatively small because patch arrangement was not manipulated directly, but selected for from essentially random landscapes. Landscape modification by humans and resulting habitat loss and fragmentation are rarely random, but follow existing vegetation patterns (e.g. preferential conversion of areas with high primary productivity) as well as jurisdictional and historical land use patterns ^5, 23^. In more structured landscapes, network properties should influence regional adaptation even more.

Network and graph theoretic perspectives on landscape connectivity provide a valuable tool for the representation and analysis of habitat patch arrangement in landscapes ^24^. Where spatial planning necessitates the prioritisation of one area over another, network measures may guide the decision so as to maintain connectivity between increasingly isolated patches ^25, 26^. Removal of patches with high betweenness-centrality, for example, has been shown to reduce connectivity and impair spatial insurance effects that maintain metacommunity robustness to habitat loss ^27^. Preservation of patch connectivity over larger regions is particularly relevant in the light of climate change that may necessitate range shifts for species persistence ^28^. Because we here focused on whole-metacommunity response capacity, we limited the analysis to network-wide metrics.

In our models, higher levels of clustering were beneficial for response capacity. In clusters of better connected patches (components) higher response diversity may accumulate as long as patches are sufficiently different in environmental conditions ^27^. Provided that occasional dispersal events could connect clusters, this was beneficial for the whole region. In a metapopulation framework, higher clustering or aggregation of patches has been found to slow down the speed of advance of range shifting species but increase the proportion of patches occupied ^29^, allowing for higher degrees of species persistence.

The characteristic path length of a network is the average of all shortest paths between any pair of nodes. Shorter average path length indicates good overall patch reachability and/or the presence of patches that form stepping stones across the landscape, facilitating traversal and range shifts, and improving regional response capacity. Corridors of stepping stones facilitate and accelerate species range shifts across larger spatio-temporal scales ^28–30^.

High clustering and short characteristic path lengths appeared to allow for alternative response mechanisms, such that either high overall reachability but low clustering, or low characteristic path lengths but high clustering could maintain response capacity (Suppl Fig S2).

An intriguing result was that a larger leading eigenvalue of the connectivity network corresponded to higher response capacity but only at the greatest inter-patch distances, where successful dispersal was rare and highly stochastic. In classic metapopulation theory, the leading eigenvalue of an analogous matrix of colonisation probabilities has been termed the metapopulation capacity of the landscape ^31^. As long as the leading eigenvalue exceeds a threshold value determined by species properties (the ratio of extinction to colonisation rate), there is long-term species persistence in the landscape. How this relates to the multispecies metacommunity dynamics modelled here is not entirely clear. The dominant eigenvalue appears to be related to percolation through sparse networks ^32^, and thus could be relevant to probabilities of random spread through the network when patches are scarce. The relevance of the dominant eigenvalue of the connectivity network in metacommunities clearly deserves more detailed investigation than we are capable of providing here.

### Implications for ecosystem functioning

Shifting climatic niches not only pose problems to individual species, but affect whole community functioning. Trait-lags of mean community performance traits directly translate to changes in rates of primary productivity. A larger trait-lag corresponds to a situation where more species in a community have suboptimal trait values, such that productivity rates are lower. Our results demonstrate effects of climate change on regional productivity that persist long after climate change has come to a halt. Habitat fragmentation that limits dispersal thus not only threatens individual species, but also leaves a legacy of impeded community functioning (a functioning debt ^20^).

One advantage of a focus on trait distributions is that they can represent a direct link between species responses and species effects on ecosystem properties ^3, 33^. A recent analysis of community responses to a long-term fertilisation treatment for example showed a directional shift in a mean community trait (specific leaf area, often correlated with individual growth rate ^34^) that tracked the continuously changing optimum, along with decreases in trait variance and increases in net primary productivity ^3^. The temperature optimum trait modelled here likely depends on multiple, co-varying traits that influence growth, i.e. fitness depends on the adaptive value of the multidimensional, integrated phenotype ^35^. Climate change further is comprised of changes in precipitation and other variables as well. Next steps in trait-based ecology should focus on establishing links between multiple traits and species vital rates or fitness components in different environments ^36^, as well as effects of these traits on ecosystem processes other than primary productivity. Trait-based models as the one presented here can incorporate multiple co-varying traits ^13^, perhaps even also traits that mediate the elusive dispersal capacity of species ^37, 38^. A comprehensive understanding of trait covariation would allow predictions regarding changes in composition and structure of plant communities and concurrent impacts on multiple ecosystem processes in response to climate change and habitat fragmentation.

### Limitations

As described here, our modelling approach has a number of limitations that could be amended by altering the setup. To disentangle effects of dispersal rate, environmental fluctuations and patch arrangement, the model could be changed as follows. First, connectivity networks could be manipulated directly via patch arrangement to create stronger gradients of network property values ^39^. Alternatively, the connectivity networks of real landscapes could be derived from one or more given dispersal kernels to study the range of network metrics observed in real landscapes along gradients of human influence. Second, dispersal rate could be manipulated directly, e.g. by altering biomass allocation to seeds. Third, inter-annual temperature variability could be either fully de-synchronised, such that patches fluctuate independently, or the degree of synchronisation could decay with spatial distance. A comparison with scenarios with no variability would inform on the magnitude of the stabilising effect of fluctuations ^40^. Velocity and magnitude of the warming curve could also be manipulated to study effects of different climate change scenarios. Fourth, forms of competition other than the generalised Lotka-Volterra formulation could be considered. Introducing resource competition, for example, would allow to introduce a second habitat quality factor (resource availability), such that the interplay of two traits (temperature optimum and resource uptake rate) could be studied. Lastly, a simplified setup would relax computational and data storage constraints and allow to track also local trait distributions and their variability within the landscape. To better assess commonalities and differences with other models, local and regional diversity measures could be recorded.

### Conclusions

Trait distributions are a powerful and versatile conceptual tool to study community responses to climate change. Recasting biodiversity theory in terms of traits integrates the mechanisms that generate and maintain diversity and allows scaling from individual fitness through community trait distributions to ecosystem properties ^3^. We have applied theory that predicts a trait-lag in response to directional environmental change ^10, 13^ to the metacommunity level to propose a simple metric of integrated regional response capacity, the inverse of the integrated lag. Using this metric, we have demonstrated how local compensation from resident species combines with immigration from the region to reduce the trait-lag and maintain metacommunity functioning under changing conditions. Species range shifts were crucial for the maintenance of functioning and connectivity network properties had a noticeable effect in larger landscapes. Yet trait-lags could persist throughout and beyond climate change, implying long-lasting reductions in meta-community functioning.

## Methods

We developed a spatially explicit metacommunity model of perennial plants set in a fragmented landscape characterised by a latitudinal temperature gradient (1°C / 100km). All 100 patches were populated with 100 species at equal initial abundances. Species had different trait values of *T*_*opt,i*_, which denoted their thermal niche optimum and determined temperature-dependent growth rates according to the Gaussian function

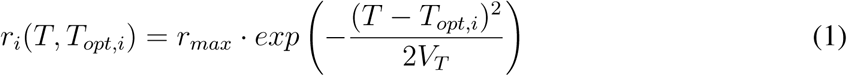

where *r*_*max*_ is equal for all species; *T* is the current ambient temperature; *V*_*T*_ is the variance of the temperature response curve (see Suppl Table S1 for parameter values). The model followed the the mean and variance of the local biomass weighted trait distribution of *T*_*opt*_ over time *t* averaged across all patches.

Growth during each growing season (180 days per year) was modelled as Lotka-Volterra competition

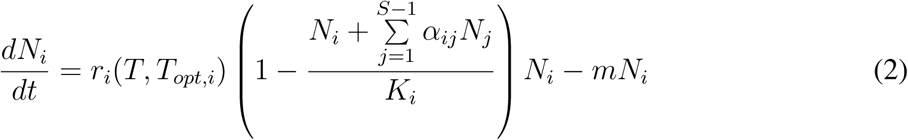

where *S* is the total number of species in a community; carrying capacity *K*_*i*_ = 1 for convenience; the interspecific competition coefficient *α*_*ij*_ < 1, ensuring coexistence; *m* is background mortality.

After each growing season, species allocated a fixed fraction (Fecundity) of their realised local biomass to seed production. Seeds dispersed according to a Poisson random draw from the 2D dispersal probability distribution (kernel)

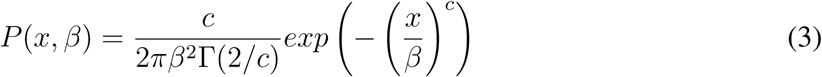

where *x* is distance; *c* = 0.5 (leptokurtic); *β* determines mean dispersal distance. Success of dispersal events were stochastic, hence actual functional connectivity was not fully determined. For each propagule parcel sent out by a species, a random number was drawn from a Poisson distribution generated with *λ* based on the mass of propagules in the parcel. The larger the propagule parcel, the more likely it arrived successfully.

A fixed proportion of biomass overwintered locally and formed, together with immigration from seed dispersal, each species’ initial abundance *N*_*i*_ the next year.

To avoid edge effects, landscapes are wrapped in East-West direction (i.e. right and left edges were joined), and extended in North-South direction with copies of the central landscape. Abundance distributions in communities of this surrounding region are extrapolated from community *T*_*opt*_ distributions within the landscape by linear regression from observed CWM *T*_*opt*_(*x*) as a function of latitude. The species pool’s *T*_*opt*_ distribution is assumed to be normal, with a variance corresponding to the mean variance of communities within the landscape as a function of time.

The model was run for 200 years to allow communities to reach local equilibrium. Then, climate change (CC) started and temperatures increased uniformly following a sigmoid warming curve that saturated at *CC*_*max*_ = 5°C above current temperatures in each patch, with a half-saturation time of 75 years. CC followed the upper range of the intermediate IPCC emission scenario RCP 6.0 ^41^.

Environmental temperature *T* exhibited temporal heterogeneity between years across the whole landscape. In all patches the same time series of autoregressive Gaussian noise was added to *T*, generated with an AR(1) model of environmental noise as *y*_*t*_ = *ϕy*_*t*−1_ + *ϵ*_*t*_, where *ϕ*_1_ = 0.6 and *ϵ*_*t*_ is an uncorrelated Gaussian innovation process with mean zero and variance 0.02 (range ≈ {−0.9 < *y* < +0.9}). Temperature noise affected local species dynamics, and we therefore replicated the whole simulation set five times with different randomly generated autoregressive noise series.

### Setup

Keeping the number of patches constant, we manipulated the extent of the landscape and therewith inter-patch distance (5 levels: 1, 2, 4, 8, 16 km of average nearest neighbour distance) as well as connectivity. Landscapes were characterised by properties of their functional connectivity network, calculated from patch arrangement and the dispersal kernel (without stochasticity). Network properties used were characteristic path length (lambda, calculated with the MATLAB BC toolbox ^42^)), the leading eigenvalue of the connectivity matrix, and a measure of spatial clustering based on Morisita’s index ^43^, calculated based on the *miplot* algorithm in the R package *spatstat* ^44^. For each inter-patch distance, we first generated 100’000 random patch-distributions. We then selected landscapes for a full factorial combination of high and low values for each of the network properties (2^3^ = 8 levels), with 5 replicate landscapes for each combination. Network values were normalised for analysis to make them comparable between landscapes of different extents.

Additionally, interspecific competition *α*_*ij*_ (0.5, 0.8) and maximum growth rate *r*_*max*_ (0.01, 0.1) were manipulated in a full-factorial design (levels in brackets).

### Partitioning of local and regional components

Each parameter combination was run with three dispersal scenarios, using the same landscape and noise series: No dispersal (SS: Species Sorting), full dispersal (Disp), and one-step dispersal or no-traversal (NoTrav). In the latter (NoTrav) we generated a look-up table for each site in the landscape as to whether a species had been present in the site during the 25 years before climate change begins (year 200). Only species present were allowed to disperse further; species that had not been present were allowed to immigrate but further traversal was prohibited. This allowed us to separate the dispersal component arising from species traversing the landscape using patches as stepping stones.

### Estimating time-scales

For each simulation and each site the trait-lag (TL) over time was calculated as the difference between CWM *T*_*opt*_(*t*) and current ambient temperature. TL was then averaged across all sites to provide one time series of the regional average trait-lag TL_MC_ for each simulation.

The trait-lag for the simulation without any dispersal shows the capacity of community response by local sorting of species present at the start of climate change (*TL*_*SS*_). The trait-lag for the simulation without any dispersal subtracted by the trait-lag of the simulation with full dispersal shows the partial capacity of community response attributed to dispersal (*TL*_*Disp*_). The trait-lag for the simulation with full dispersal subtracted by the trait-lag of the simulation with only nearest site-neighbour dispersal shows the partial capacity of community response attributed to traversal of species through the landscape (*TL*_*Trav*_).

We then fitted a function to these trait-lag components as

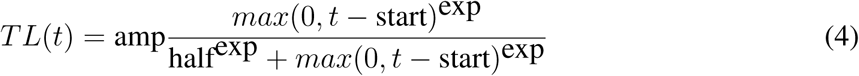

parametrized by four variables, the maximum amplitude (*amp*), the delay until the signal induced by climate change begins (*start*), the time until half of the maximum effect is reached (*half*) as well as the shape of the function (*exp*) which for all values > 0 is saturating and for values > 1 can have different degree of sigmoidal shape. Note that climate change was applied as a function with *amp* = 5, *start* = 200, *half* = 75 and *exp* = 3. Thus, for each potential part of the landscape-level response (SS, Disp and Trav) we could determine the size of the effect (*amp*), the time scale (*start* and *half*) as well as the shape of the dynamics.

Combining these component functions yielded a near perfect fit to the time dynamic of the overall regional trait-lag TL_MC_ such that *TL*_*MC*_(*t*) = *TL*_*SS*_(*t*) − *TL*_*Disp*_(*t*). This enabled us to summarise the whole landscape response by integrating TL_MC_ over time, and we thus could define response capacity as 1/ TL_MC_. The time dynamic of TL_MC_ was further characterised by recording the maximum trait-lag experienced (**TL**_**Max**_), the halftime of the initial development of TL_Max_ (**t**_**Dev**_) and the time it took for the trait-lag to recover to half of the maximum (**t**_**Rec**_).

### Analysis

A factorial ANOVA was conducted to compare main effects and interactions of the four factors inter-patch distance (5 levels), *α*_*ij*_ (2 levels; weak 0.5, strong 0.8), *r*_*max*_ (2 levels: slow 0.01, fast 0.1) and network combination (2^3^ = 8 levels: all possible combinations of Low and High values of the three network properties) on response capacity. Rather than assessing statistical significance, we were interested in effect sizes, reported as *η*^2^. Analyses were performed in R version 3.3.0 ^45^.

The model was written in Julia language ^46^.

## Acknowledgements

This research was performed under the Strategic Research Program EkoKlim, Stock-holm University.

---

**Supplementary Figure S1:**
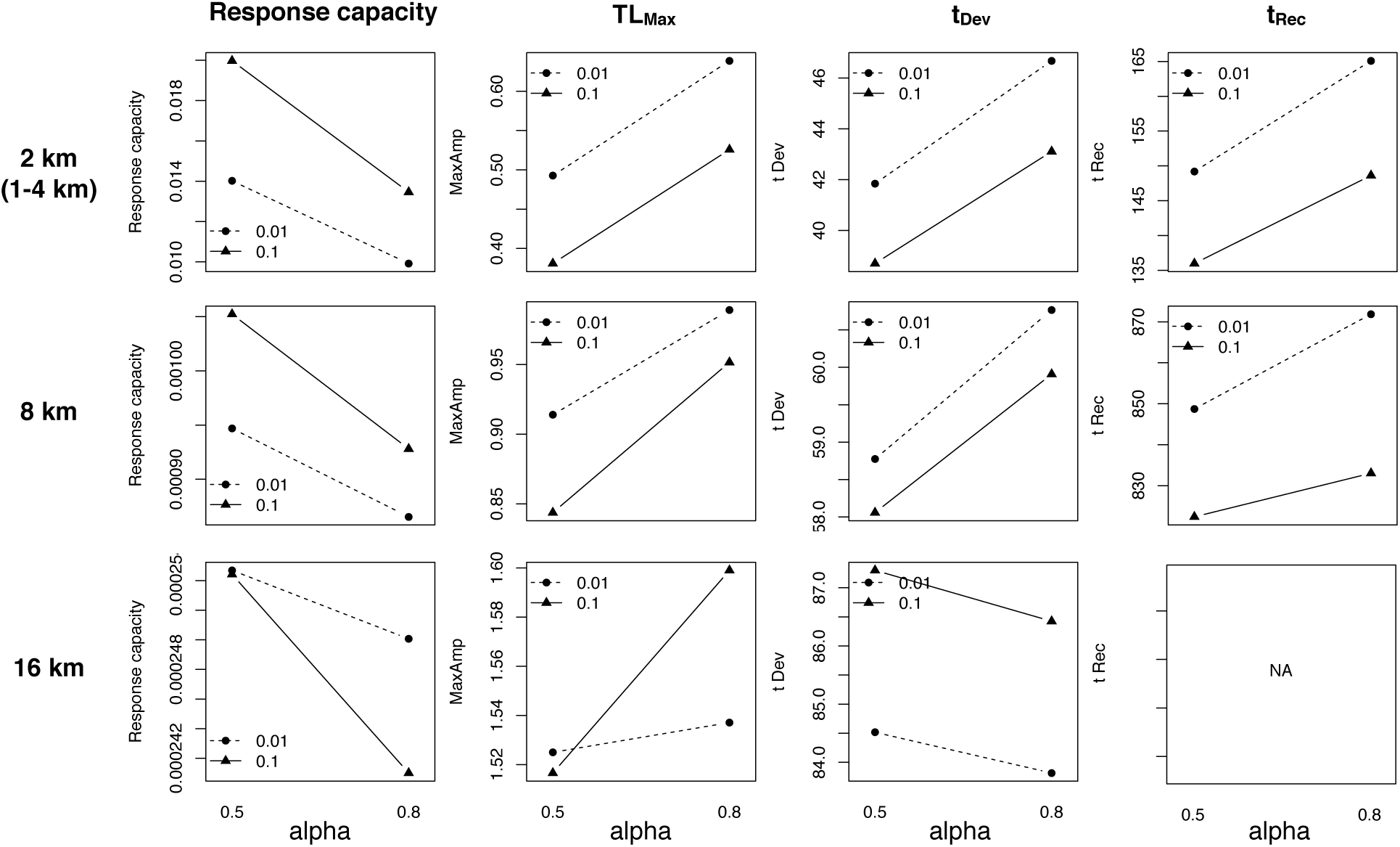
Interaction plots of mean response capacity, the maximum trait-lag TL_Max_ and timepoints of maximum trait-lag t_Max_ and recovery to half of TL_Max_, t_Rec_, at different patch distances, different levels of interspecific competition (alpha) and maximum growth rates (trace: dashed line: *r*_*max*_=0.01, full line: *r*_*max*_=0.1). Intermediate patch distances from 1-4 km showed very similar responses and are here represented by figures for distance = 2 km. Response capacity and TL_Max_ responded similarly to factor levels: metacommunities with higher response capacity had a low TL_Max_. Responses of timescales changed with extent, except for intermediate distances of 1-4 km. Note the flip at distance = 16 km for all variables. t_Rec_ was not reached at distance = 16 km.

**Supplementary Figure S2:**
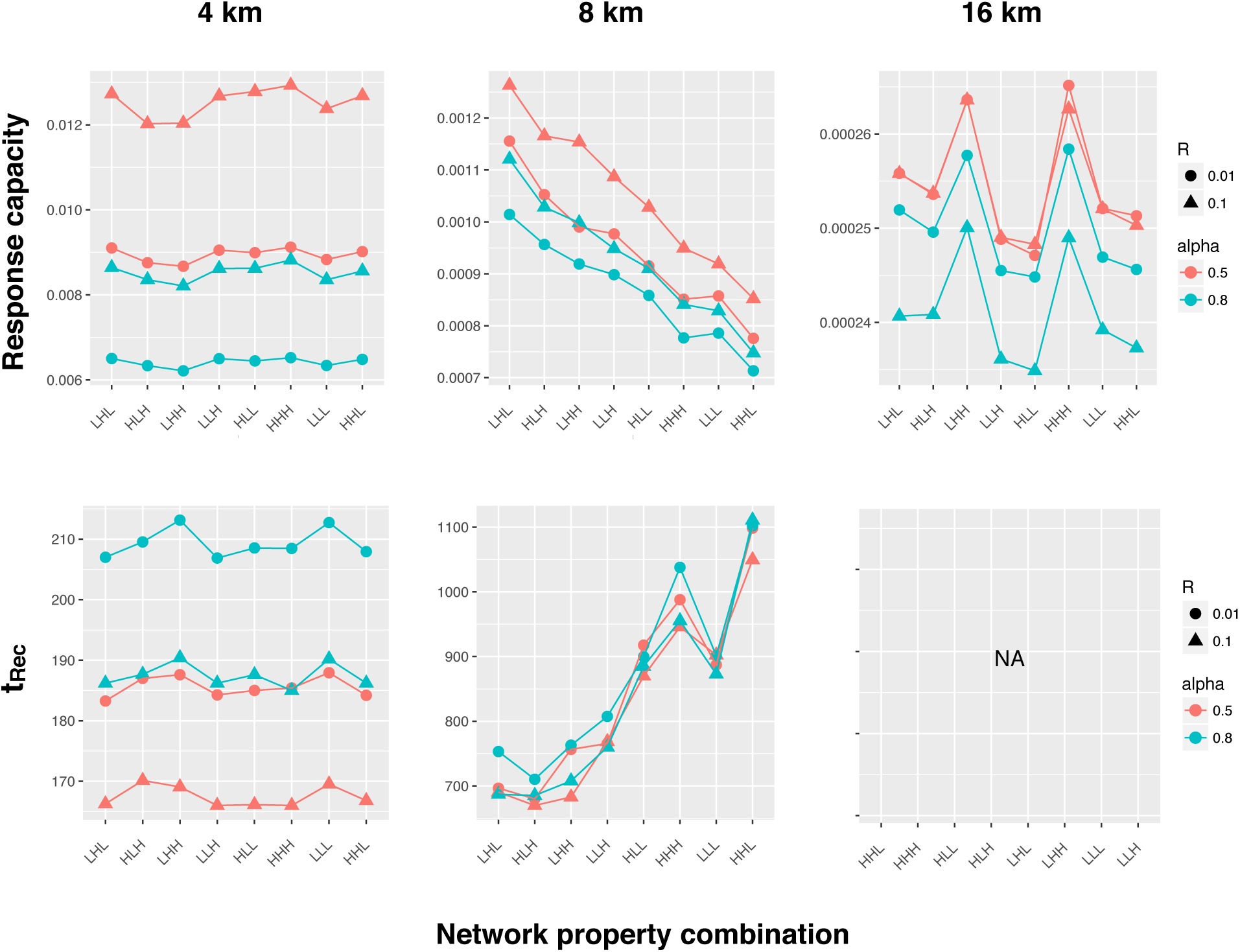
Effect of network property combinations on recovery time t_Rec_ and response capacity for the three largest landscapes (inter-patch distance levels 4, 8, 16 km). Group levels refer to combinations of low (L) and high (H) values of the 3 network properties: characteristic path length (lambda), the leading eigenvalue, and the clustering index. Network combination levels are sorted for response capacity at distance 8 km. At shorter inter-patch distances (not shown), response capacity did not vary with network combination.

**Supplementary Figure S3:**
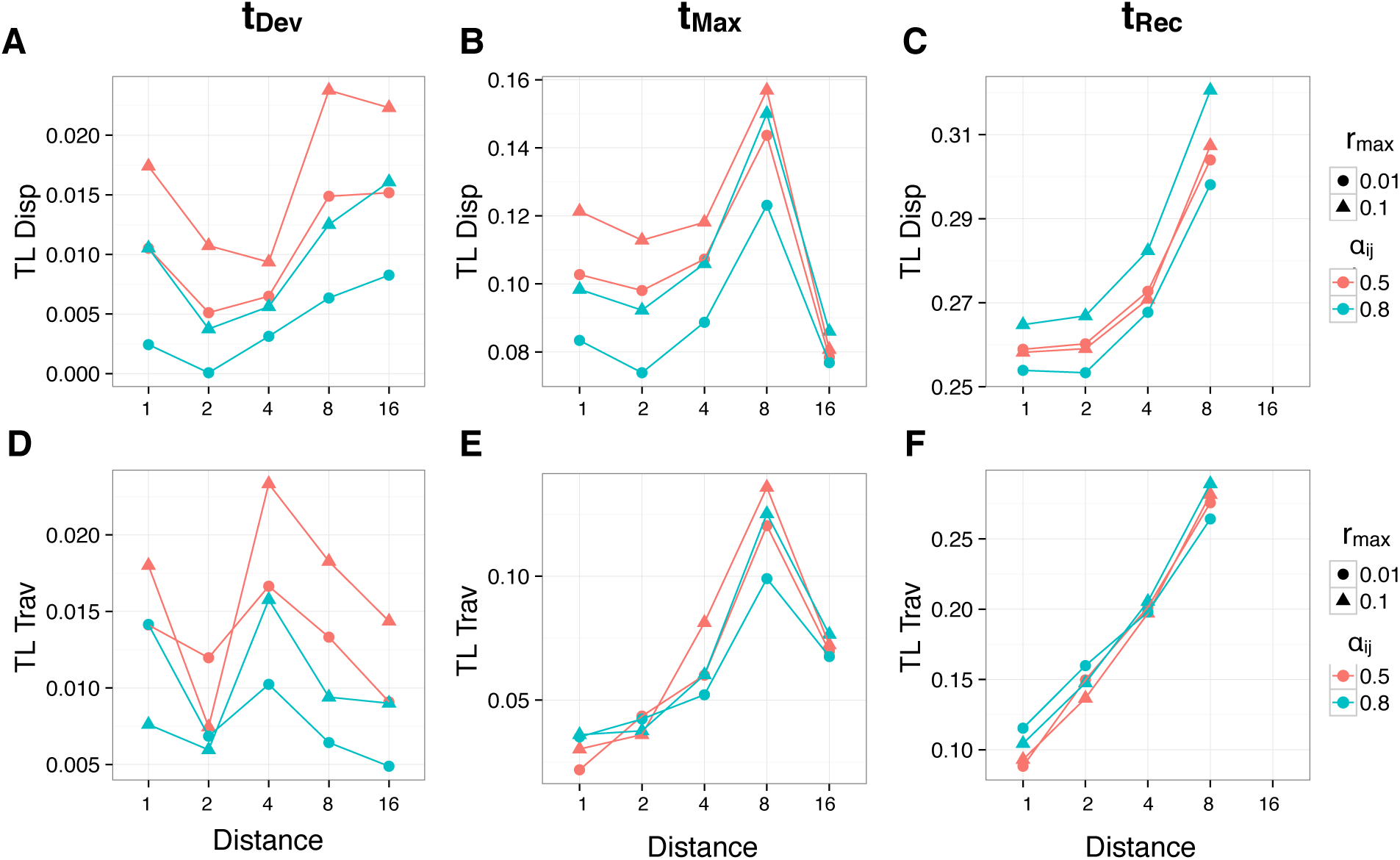
Proportional contribution of components to lag reduction at t_Dev_, t_Max_, and t_Rec_. A-C: Proportional contribution to lag reduction by dispersal (TL_Disp_, including traversal). The remaining proportion of lag reduction (1-TL_Disp_) was due to initial local species sorting of resident species (not shown). D-F: Proportional contribution to lag reduction by traversal (TL_Trav_, part of dispersal).

**Supplementary Figure S4:**
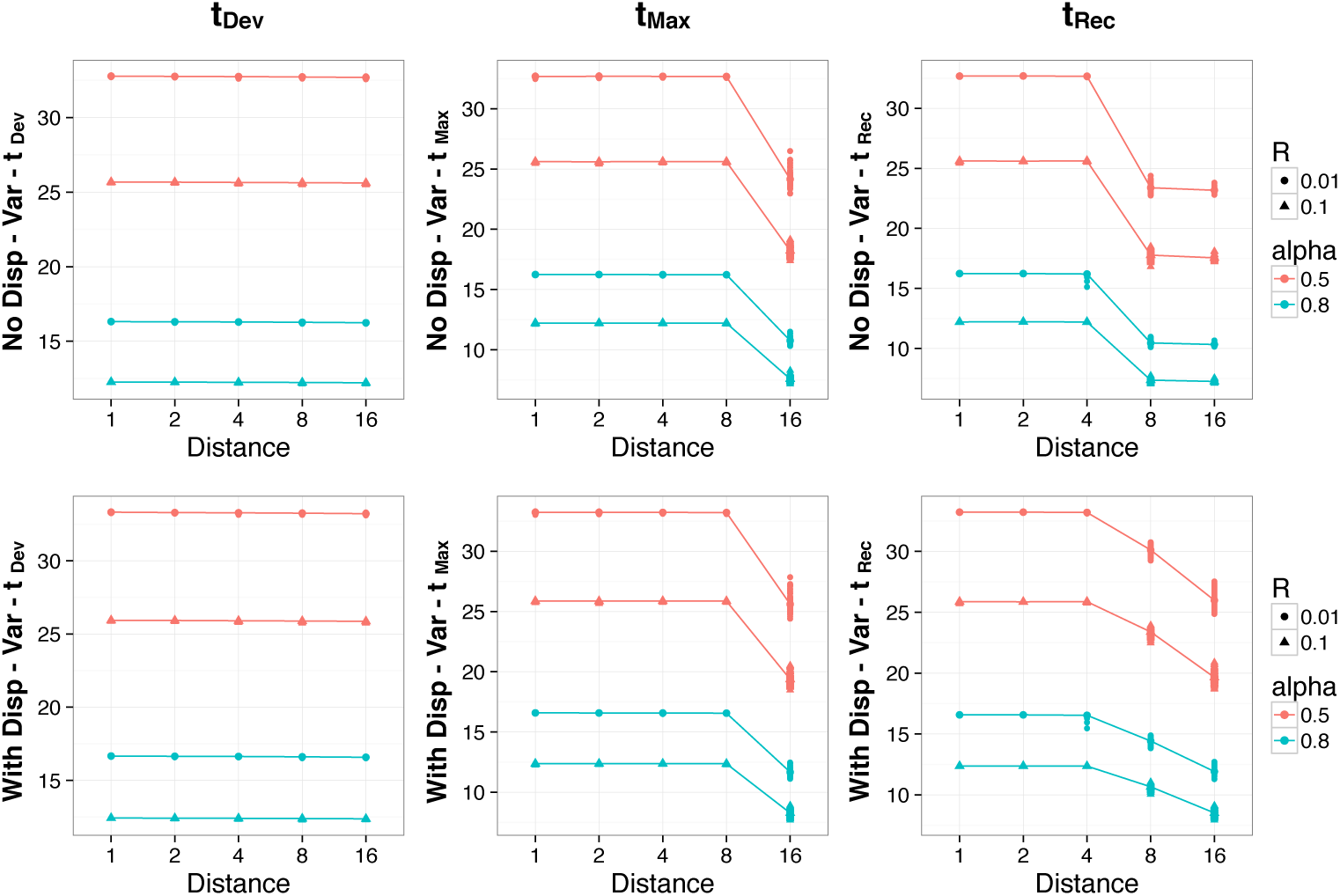
Average variances of community trait distributions at three timepoints (t_Dev_, t_Max_, t_Rec_) for different factor levels, with (top row) and without (bottom row) dispersal.

**Supplementary table S1:** Model parameters overview.

| Parameter | Values | Comment |
| --- | --- | --- |
| **Manipulated** |  |  |
| $\alpha_{ij}$ | 0.5, 0.8 | Interspecific competition |
| $r_{max}$ | 0.01, 0.1 | Maximum growth rate |
| Extent | 10, 20, 40, 80, 160, 320 | Side of square landscape |
| **Fixed** |  |  |
| Mortality $m$ | $0.05 \cdot r_{max}$ | Background mortality |
| $V_T$ | 45 (90) | Variance = 45 |
| Range of T<sub>opt</sub> | 0 to 40 °C | Evenly spaced (diff = 0.4) |
| No Species | 100 |  |
| Fecundity | 0.05 | Allocation to seed production |
| Carrying capacity K | 1 |  |
| Extinction threshold | 1.0e-10 | One seed per K |
| Overwintering fraction | 0.3 | Perennials |
| CC max Temp increase | 5 °C |  |
| CC halftime | 75 yrs |  |
| Dispersal kernel: $c$ | 0.5 | Leptokurtic |
| Dispersal kernel: $\beta$ | 20 | Mean distance $\delta = 0.4\text{km}$ |
| AR noise time series Var | 0.02 | Range $\approx \{-0.9 < y < +0.9\}$ |
| Temp gradient | 1 °C/ 100km | Landscape centre: 20 °C |
| No Sites | 100 |  |

